# Shear Stress Conditioning Promotes a Pro-Inflammatory Response in Porcine Endocardial Endothelial Cells

**DOI:** 10.1101/2025.02.03.636291

**Authors:** Pengfei Ji, Kathryn Jane Grande-Allen, Swathi Balaji, Ravi K. Birla, Sundeep G. Keswani

## Abstract

**PURPOSE:** Discrete subaortic stenosis (DSS) is a congenital heart disease characterized by a narrowing of the passage below the aortic valve in the left ventricular outflow tract (LVOT) [1]. While endocardial endothelial cells (EECs) are known to play a role in DSS, the response of these cells to shear stress is not known. In this study, we hypothesize that the response of EECs to shear stress in the LVOT is a mediator of DSS.

**METHODS:** To test this hypothesis, we conditioned porcine EECs to controlled shear stress regimes using cone and plate bioreactors. Subsequently, we quantified the concentration of proinflammatory cytokine in the conditioned media using the Luminex assay. Bulk-RNA sequencing was used to quantify changes in the genotype of the shear stress conditioned EECs.

**RESULTS:** The expression of CD31 was knocked down and subsequently, the changes in release of shear stress induced proinflammatory cytokines released by EECs quantified using the Luminex assay. The results of these studies show that the inflammatory cytokines were highly selected in the conditioning medium, and under bioreactor treatment the cell activated the PI3K-AKT and TNF-a signaling, which also triggered the other immune cell responses though Th1, Th2 and Th17 cell differentiation pathways. Furthermore, CD31 was identified as a mediator of the pro-inflammatory response of shear stress conditioned EECs.

**CONCLUSIONS:** The studies provide a clear link between shear stress, and the subsequent proinflammatory response of EECs as a mediator of DSS.

## INTRODUCTION

Discrete subaortic stenosis (DSS) is a congenital heart disease caused by the formation of a fibro-membranous tissue, which results in increased blood pressure in the left ventricular outflow tract (LVOT) [1]. Currently, the main treatment method to manage DSS patients is surgical resection of the fibrotic membrane. However, there are risks associated with the surgery including significant risks associated with heart bypass, sternotomy, and anesthesia, and the high incidence of reoccurrence, which requires reoperation [2]. The main problem with the management of DSS patients is the reoccurrence of the fibrotic membrane, which requires repeat surgery.

There are some hypotheses for the pathogenesis of DSS, and one mainstream opinion from genetic etiology is that the correlation between DSS and left ventricular (LV) geometric abnormalities has provided strong evidence for a hemodynamic cause. This suggests that congenital or post-surgical LVOT geometric irregularities generate anomalous shear forces on the septal wall, leading to a fibrotic response [3]. The etiology for DSS and potential mechanical cell signaling pathway resulting from the shear stress response of endocardial endothelial cells (EECs) in the LVOT are still unknown and remain a knowledge gap. Furthermore, the subsequent effect on fibroblasts leading to the formation of the fibrotic membrane in DSS remains unknown. Therefore, in this study our goal was to characterize the effect of shear stress on EEC and the subsequent response of these cells based on cytokine release and changes in the gene expression profile based on bulk RNA sequencing.

Mechanobiological experiments are the ideal methods to ensure unidirectional blood flow between the LV and the aorta. These studies involve exposing cells to a mechanically demanding environment that involves various stresses such as bending and tensile stress, pressure, and fluid shear stress [4]. Thus, motivated by the design of a bioreactor that can alter the fluid and shear stress force, the cone-and-plate geometry has been described and can influence the oscillatory flow. In addition, the flow is analyzed by two parameters: one is the Womersley number and the second is the Reynolds number. Both parameters are determined by the local acceleration and centripetal which was influenced by the speed and diameter [5]. Specifically, the mechanisms of the function of heart valves is determined by the surrounding hemodynamic environment, and the interactions between heart valve and fluid dynamics are vital to understanding the heart valve function and DSS [6]. As described above, the cone-and-plate device produces a flow with shear stress on the surface of the plate as quantified by the following equation. T=pvw[r/(h+ra)] [5]. The cone-and-plate bioreactor (CPB) is a device to investigate the heart valves, the pathology and the potential mechano-etiology of the DSS [7].

In this study, the CPBs were used to understand the influence of altered shear stress in the LVOT of DSS patients on EECs. The CPB has an inverted cone rotating above a flat stationary plate. Fluid flows within a gap formed between the cone and the plate. This process distributes uniform shear stress on the surface [8]. The magnetic stir plate adjusts the cone’s angular velocity by modifying the RPM on the control unit and subsequently, this allows control over the magnitude of the shear stress applied to the EECs. We defined 4 different shear stress regimes for this study: negative control, positive control, low shear stress and high shear stress, based on our published computational models for DSS [8]. Negative control = 0 dynes/cm^2^ or 0 RPM, positive control = 6 dynes/cm^2^ or 100 RPM, Low shear stress = 15 dynes/cm^2^ or 280 RPM and high = 35 dynes/cm^2^ or 560 RPM.

The endocardium is the innermost layer of the heart wall, which is formed by the endocardial cells, and endocardial cells are one kind of endothelial cell [9]. Genetic techniques such as RNA-sequence and genetic lineage-tracking technology revealed several unexpected roles of the endocardium during heart development [10]. In particular, the endocardial cells produced a physical barrier against the cardiac chamber, and it is also an important cell source to form a functional heart [11]. Recent studies showed that endothelial-to-mesenchymal transition (Endo MT) in the heart can also give rise to diverse cell lineages beyond cushion mesenchymal cells. For example, Endo MT-derived cells have been shown to differentiate into cardiac fibroblasts, smooth muscle cells, pericytes, and endothelial cells. These cells play important roles in the maintenance of cardiac structure and function by using genetic lineage-tracing technology [12]. However, the abnormal formation of the endocardial layer between endocardium and myocardium leads to congenital heart diseases such as DSS, heart valve diseases and noncompaction myocardium, [13, 14, 15].

Since the lack of pediatric DSS patient heart specimens for cell isolation, we selected to use the porcine EECs as the experimental model for this study [16]. EECs were isolated from the inner layer of the LV by using a trypsin-based method, and porcine EECs showed the distinct morphological characteristics compared with HUVECs [16]. CD31, also known as PECAM-1 (Platelet Endothelial Cell Adhesion Molecule-1), is a cell adhesion molecule that is commonly used as a marker for endothelial cells. It has been reported that porcine EECs exhibited the positive expression of CD31, maintaining a CD31-positive characteristic phenotype [16]. In our study, we conducted a comprehensive characterization of differentiated ECs obtained from patients’ iPSCs and control iPSCs. Our finding indicated that iPSCs-ECs express various endothelial genes (CD31, CD144, KDR, Tie2, vWF and NOS3) through both RT-qPCR and immunostaining [17, 18].

This study focuses on porcine EECs that were recently reported as above associated with DSS. These cell fates will unveil the roles of endocardial progenitors in function, pathological progression, and potential metabolic pathways of the DSS. These studies suggest that developmental mechanisms can be conducted in promoting heart disease development and cardiac repair and regeneration. The results of this study will lead to an understanding of the factors that control endocardial plasticity and the modulation of their cellular fate conversion has the potential to reveal new therapeutic strategies for DSS.

## MATERIALS AND METHODS

### Culture of Primary Porcine EECs Culture

0.1% gelatin (Gelatin methacryloyl, bloom 300, degree of substitution 60%, Sigma-Aldrich, 900622-1G) was used to coat the surface of the tissue culture plates. 1 ml of 0.1% gelatin solution was added to a 60 mm tissue culture plate and the plate incubated at 37oC for 15 minutes. The plates were then washed three times with PBS and used for cell culture studies. Porcine EECs were cultured on gelatin coated plates. A single vial of frozen porcine EECs were recovered from a liquid nitrogen tank and placed in a 37oC water bath until the contents were 80% thawed. Porcine EECs were cultured in ECM media (Sciencell, Catalog # 1001), prepared according to manufactuer’s instructions. (Pengfei, provide details, product and catalog number here),. The EECs were plated at a density of 500,000 cells per 60 mm tissue culture plate for 2 days, at which time the cells had reached a confluency of 70% prior to bioreactor studies, as described next.

### Preparation of the Cone and Plate Bioreactors

Our system for cone and plate bioreactors (CPBs) has been described before [8]. Briefly, the system consists of the bottom of the petri dish and the cone is loaded on top. To prepare the CPB for experiments, the cones were sterilized by exposure to UV light for 3 hours. The gap height of cone was 300 μm established by placing it on a petri dish with two stacked coverslips. The collar shaft of the cone was tightened to secure the adjusted height. Subsequently, the modified cone was placed on a petri dish containing EECs and 3.0 mL of cell culture medium. The cone-and-plate components were then moved to a magnetic stir plate with an adjustable rotational speed (Cole-Parmer, #EW-84003-82). To adjust the shear stress magnitude, the desired shear stress was determined by using the provided equation [5], and this value was used to set the rotational speed of the stir plate accordingly. Thus, we utilized the 0, 100, 260 and 660 RPM as shear stress magnitudes of 0, 6, 15 and 35 dyne/cm^2^.

### Conditioning Porcine EECs using Cone and Plate Bioreactors

Porcine EECs were cultured using the CPBs using shear stress magnitudes of 0, 6, 15 and 35 dyne/cm^2^, which corresponded to RPMs of 0, 100, 280 and 660, respectively. The treatment times selected were 1, 4, 8 and 24 hours for each of the 4 shear stress magnitudes (0, 6, 15 and 35 dyne/cm^2^), and each condition was repeated in triplicates. The total number of porcine EEC samples was 48. At the end of each time point (1, 4, 8 and 24 hours), the conditioned media was collected and sampled to quantify the concentration of proinflammatory markers using the Luminex assay and the total RNA extracted from the EEC samples and processed for bulk-RNA sequencing (both of which are described in subsequent sections).

### Multiplex Luminex Assay to Quantify Proinflammatory Cytokines

MILLIPLEX® Porcine Cytokine/Chemokine 13-Plex GM-CSF (Millipore Sigma, catalog # PCYTMG-23K-13PX) enables simultaneous quantification of multiple analytes in porcine tissue/cell lysate and culture supernatant samples, as well as serum or plasma samples. The kit can measure a range of cytokines and chemokines, including GM-CSF, IFN-γ, IL-1α, IL-1ra, IL-1β, IL-2, IL-4, IL-6, IL-8, IL-10, IL-12, IL-18, and TNFα. It utilizes a 96-well format and comes with a lyophilized standard cocktail and two internal assay quality controls, allowing for accurate and reliable results. The kit can also accommodate up to 38 samples in duplicate, making it a highly efficient and effective solution for porcine cytokine and chemokine analysis [19]. The samples were from EECs conditioned medium as described earlier and a total of 48 porcine samples were analyzed by 2 Luminex assay kits to quantify the concentration of proinflammatory cytokines (GM-CSF, IFN-γ, IL-1α, IL-1ra, IL-1β, IL-2, IL-4, IL-6, IL-8, IL-10, IL-12, IL-18, and TNFα).

### RNA Isolation and Sequencing

Analyzing the transcriptome can identify the differentially expressed genes through different cell populations, also, it can help understand the gene boundary identification, variable cleavage, and transcript variation [20]. Total RNA was extracted from porcine EECs samples using the RNeasy Micro Kit (Invitrogen catalog # 12183018A), the purity was validated using the absorbance readings at A260/A280 ratio within 1.8-2.0, which was considered acceptable. Bulk-RNA sequencing and data analysis was conducted by Novogene and used to quantify differentially expressed genes, gene ontology analysis, KEGG pathway analysis and reactome pathway analysis.

### Immunofluorescence Staining of Porcine EECs

Porcine EECs were plated onto 6 cm cell plates and 3 mL of ECM added, and the cells cultured for 3 days. After that, the CPBs were used to treat the porcine EEC for an additional 24 hours. The conditioned cells were then fixed with 4% paraformaldehyde (PFA) and the PFA rinsed with PBS three times. The blocking buffer was prepared using 3% bovine serum albumin (BSA) and 1% Triton-X, and applied it to each well, followed by incubation at room temperature. The porcine EECs were stained using Anti-CD31 antibody (Abcam, #28364) diluted at 1:5 for 2 hours, and then stained with Alexa Fluor 488 for 1 hour (1:100 dilution). Subsequently, DAPI was added using a dilution of 1:5000 in PBS to each well and incubated in a dark area for 15 minutes. The antibody was then aspired and the porcine EECs washed three times using PBS and imaged using a KEYENCE BZ Microscope (BZ-X800).

### Transfection

Lipofectamine 3000 (ThermoFisher, #L3000001) was facilitated through transfection. The porcine EECs reached 40% confluency followed by a media change prior to the transfection. Mixing the Opti-MEM reduced serum medium (Gibco, #31985070) with siRNA and Lipofectamine 3000 Reagent and incubating for 15 minutes at room temperature. Continuously, adding the siRNA-lipid complex to porcine EECs and cultured for 2-4 days at 37 °C. Also, to determine the efficiency of the transfection, we utilized the pJTI™ R4 Exp CMV EmGFP pA Vector (Thermofisher, A14146) with lipofectamine 3000 as described above, which was a fluorescent positive control plasmid that can be used to estimate the transfection efficiency. Then, we confirmed that the siRNA knockdown the CD31 from the IHC result as described in the result part. After that, the CD31 knockdown EECs were cultured, and then, we repeating the bioreactor studies on the CD31 knockdown EECs. The procedures of the bioreactor treatment were described as above. Additionally, 0, 6, 15 and 35 shear stress were choosed to test the CD31 knockdown EECs for following experiments such as RNA-sequencing.

### Statistical analysis

All data was analyzed by using Unpaired two-tailed t-test, and Two-way ANOVA or one-way ANOVA with Tukey’s post hoc test. Data were presented as mean ± SEM and P value less than 0.05 was a statistical difference.

## RESULTS

### CD31 Mediates Shear Stress Response in Porcine EECs

To investigate the CD31 expression on porcine EECs, we did the immunofluorescence to unveil the porcine EECs after bioreactor treatment for 24 hours. Porcine EECs were labelled with CD31-AF488 and DAPI, and ImagJ was utilized to determine the intensity of CD31. As can been seen from Figure 1, the expression of CD31 was influenced by the magnitude of shear stress. There was no difference in the expression of CD31 between negative and positive control groups, with shear stress magnitudes of 0 and 6 dyne/cm^2^ respectively. Furthermore, as the magnitude of the shear stress was increased to 15 and 35 dyne/cm^2^, there was a subsequent increase in the expression of CD31, relative to both the positive and negative controls (**Figure 1**). In addition, the expression of CD31 was higher in porcine EECs exposed to shear stress magnitudes of 35 dyne/cm2 relative to 15 dyne/cm2 (**Figure 1**). This data clearly show the positive relationship between shear stress magnitude and CD31 expression in porcine EECs, providing evidence to support the role of CD31 as the mediator shear stress response in these cells.

**Figure 1.**
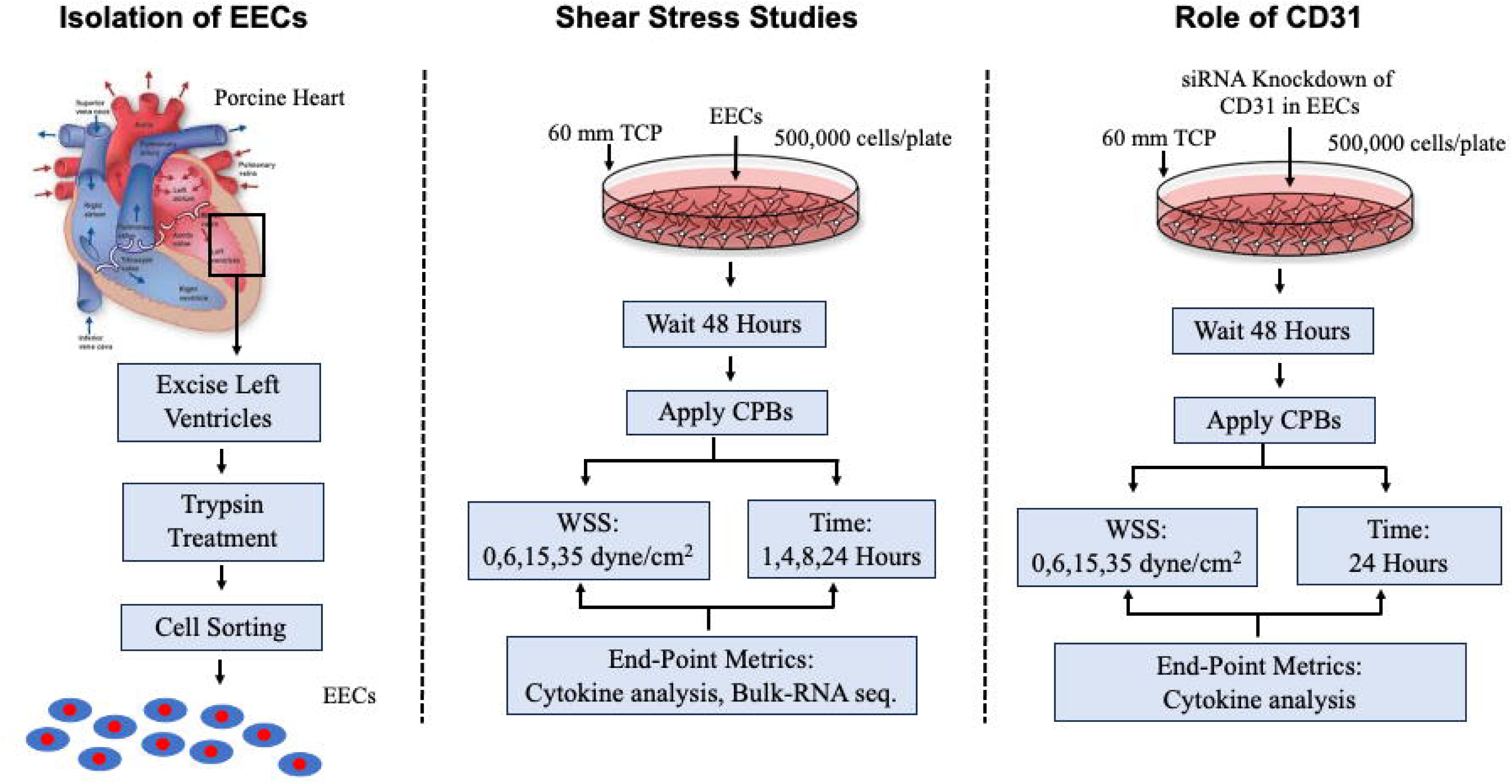
Effect of Shear Stress on CD31 Expression. Images on the left show the IHC staining as a function of time with gradually increasing shear stress (0, 6, 15 and 35 dyne/cm^2^). Image on the right shows average intensity as a function of shear stress. *p<0.05.

### Release of Proinflammatory Cytokine of Porcine EECs in Response to Shear Stress Conditioning

To further investigate the porcine EECs, we used the Multiplex Luminex Assay to analyze the composition of the porcine EEC conditioning medium in order to quantify the release of proinflammatory cytokines into the conditioned culture medium. We tested four different groups to include negative control, positive control, low shear stress and high shear stress (0, 6, 15 and 35 dyne/cm^2^) at four different time points such as 1h, 4h, 8h & 24h. The results are presented in **Figure 2**.

**Figure 2.**
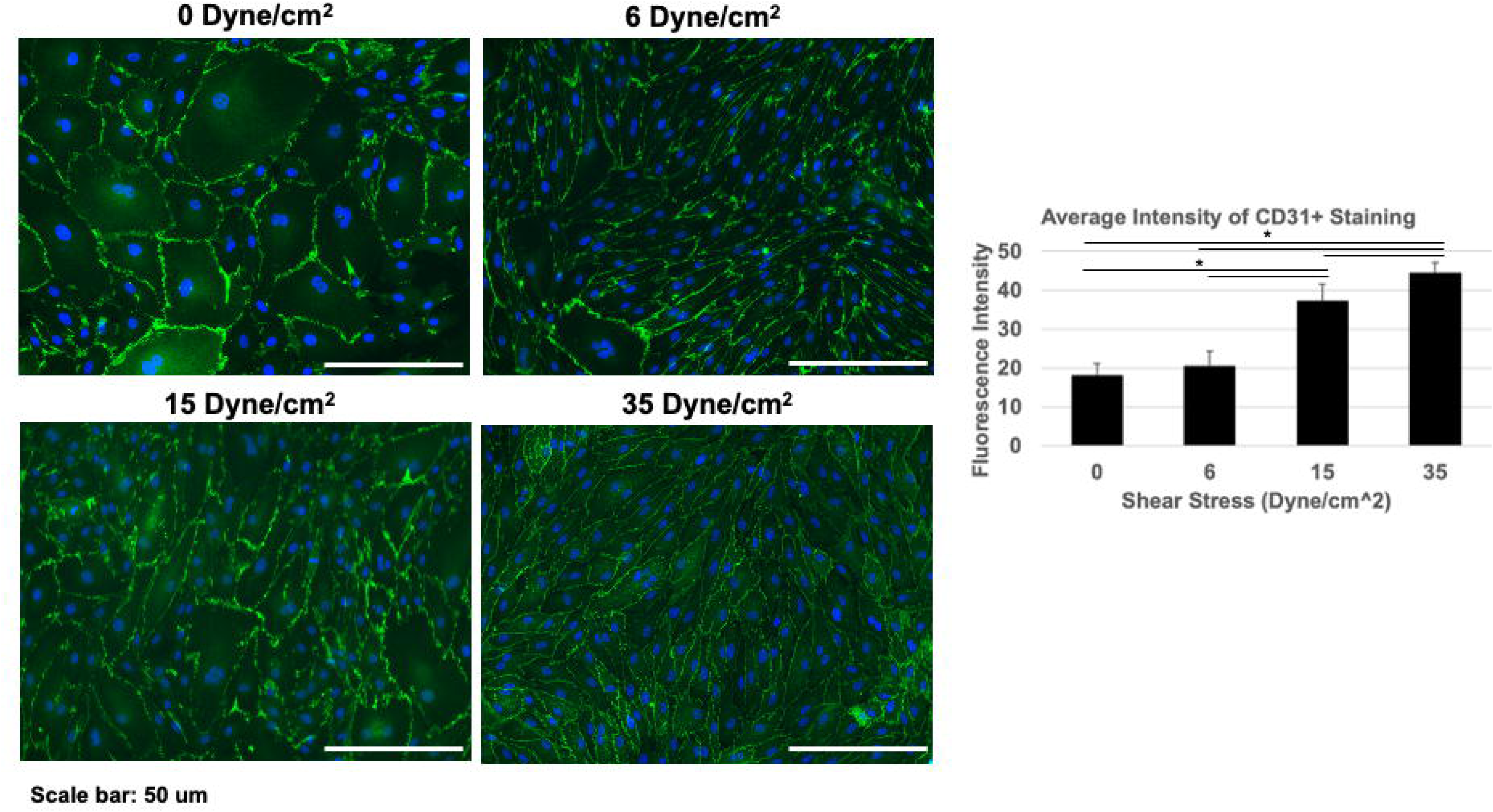
Cytokine Concentration as a Function of Shear Stress and Time. Four shear stress conditions were used and included 0, 6, 15 and 35 dyne/cm2 at four time points to include 1, 4, 8 and 24 hours. The concentration of the following thirteen cytokines was quantified using the Luminex assay: GM-CSF, IL-1b, INFg, IL-1RA, IL-1a, IL-2, IL-4, IL-10, IL-6, IL-12, IL-8, IL-18, TNFa.

At the 1-hour time point, the cytokines tend to be stable, and very few cytokines have significant differences in Figure 2. To be more specific, the several inflammatory cytokines did not alter significantly after expose to EECs to shear stress in the CPBs for 1 hour. However, significant differences were noted in the concentration of the following proinflammatory cytokines after 1-hour of expore of to shear stress in the CPBs: IFN-γ, IL-2, IL-8, IL-18 and TNFa.

A similar trend was noted after 4 and 8 hours of conditioning in the CPBs, with only a select subset of the proinflammatory significantly increasing as a function of progressively increasing shear stress magnitudes (**Figure 2**). After 4 hours, the following cytokines were found to increase: GM-CSF, IL-1b, IL-1a, IL-10, IL-6, IL-12, IL-8, IL-18 and TNFa. Similarly, after 8 hours, the following cytokines were found to increase: IL-6, IL-8, IL-18 and TNFa. After 24 hours, the following cytokines were found to increase: GM-CSF, IL-1b, IL-1RA, IL-1a, IL-2, IL-6, IL-10, IL-12, IL-18, and TNFa.

Collectively, the data presented in **Figure 2** serve to demonstrate a progressive increase in the proinflammatory response of porcine EECs as a function of shear stress magnitude and time, with different cytokines peaking at different time points.

### The Proinflammatory Response of Porcine EECs to Shear Stress Confirmed using Bulk-RNA Sequencing

To detect the porcine EEC cell signaling pathway after bioreactor treatment, we conducted bulk-RNA-sequencing to identify differentially regulated genes and pathways responsible for the shear stress mediated response of porcine EECs (**Figures 3, 4 and 5)**. **Figure 3** shows the differentially regulated genes for all 24 groups, presented in the form of a volcano plot, followed by tables listing the top 10 most significant genes both upregulated and downregulated for each of the 24 groups. Finally, the total number of differentially expressed genes is shown as a bar graph, differentiating both upregulated and downregulated genes [21].

**Figure 3.**
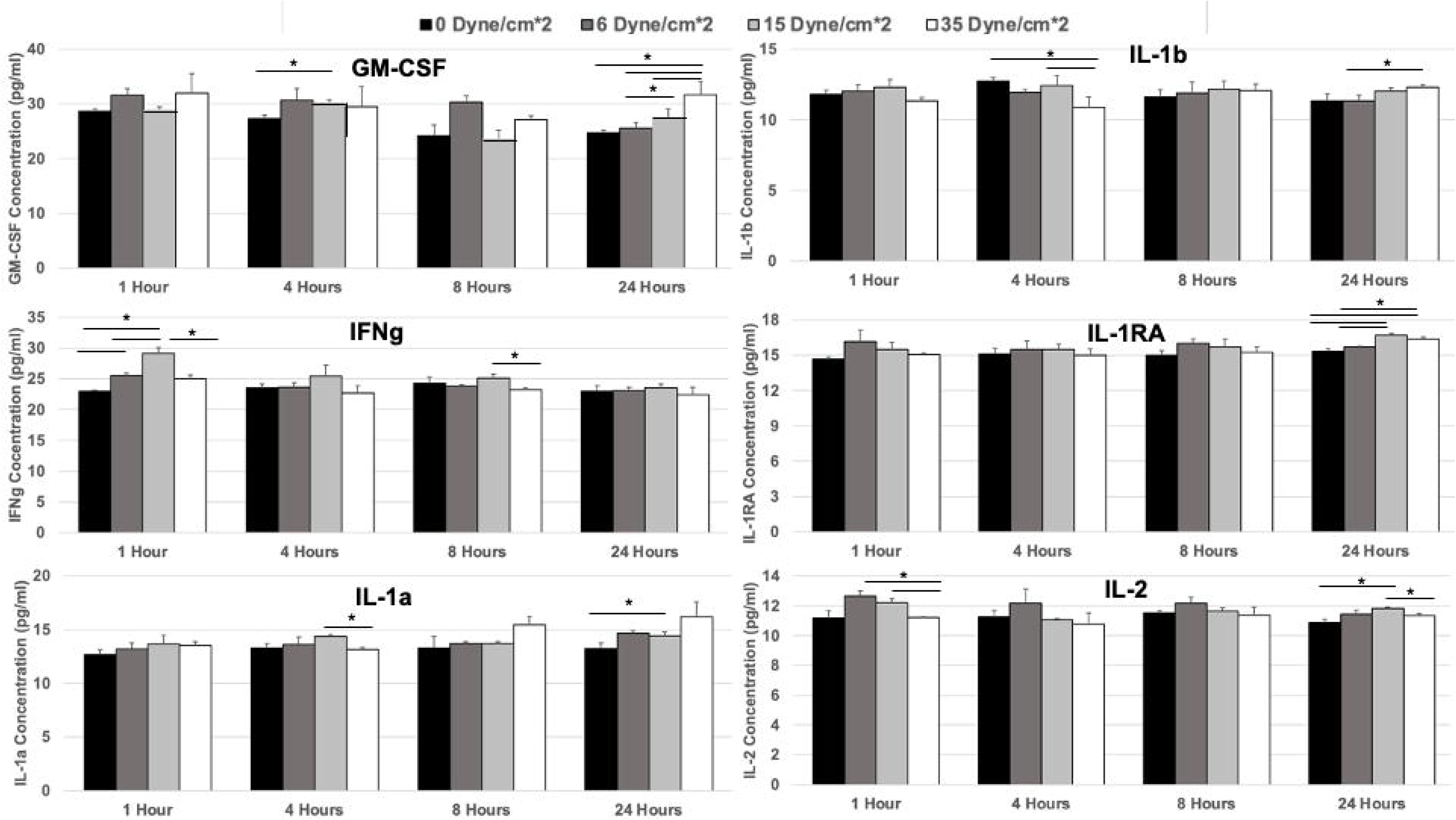
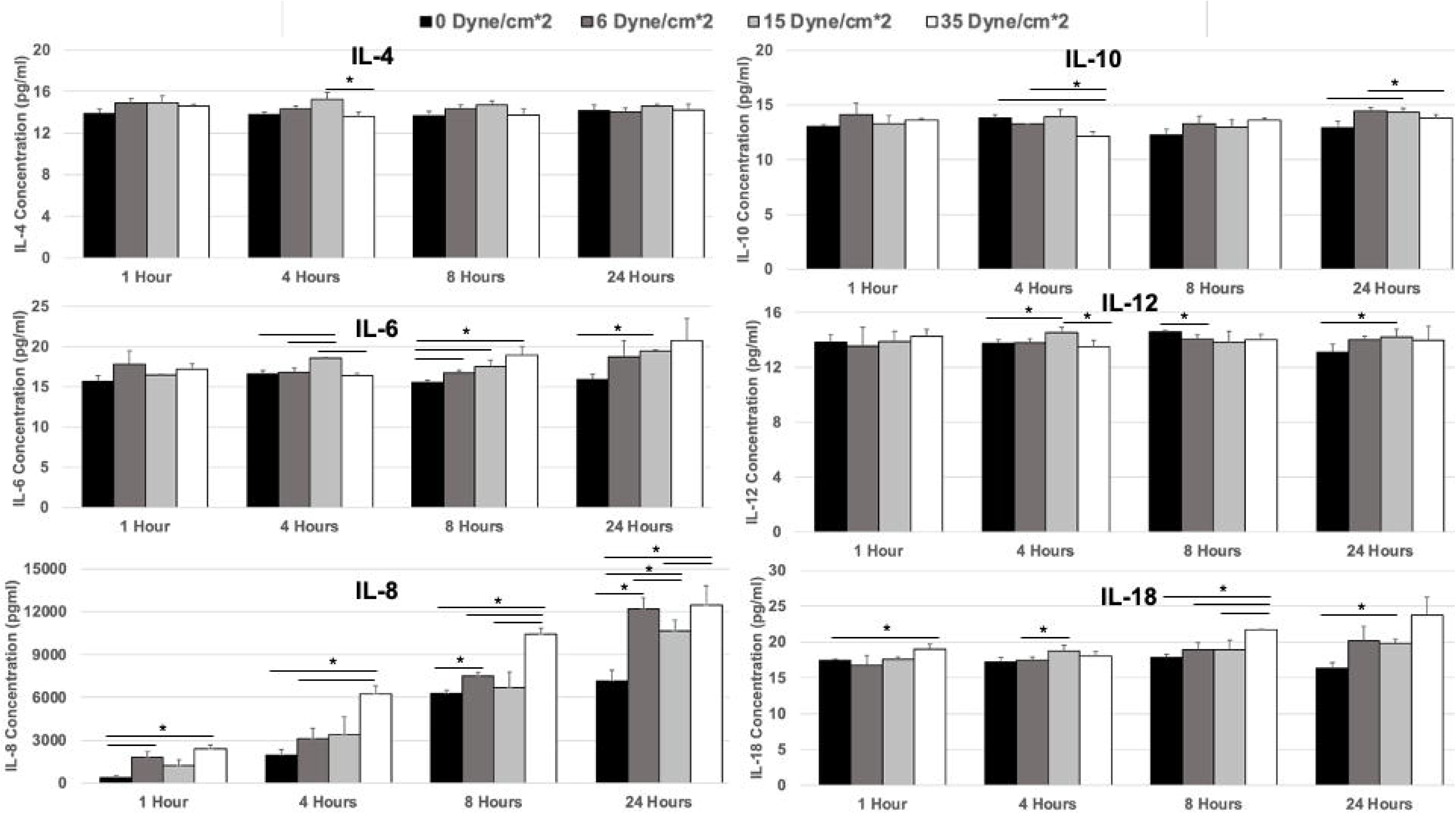
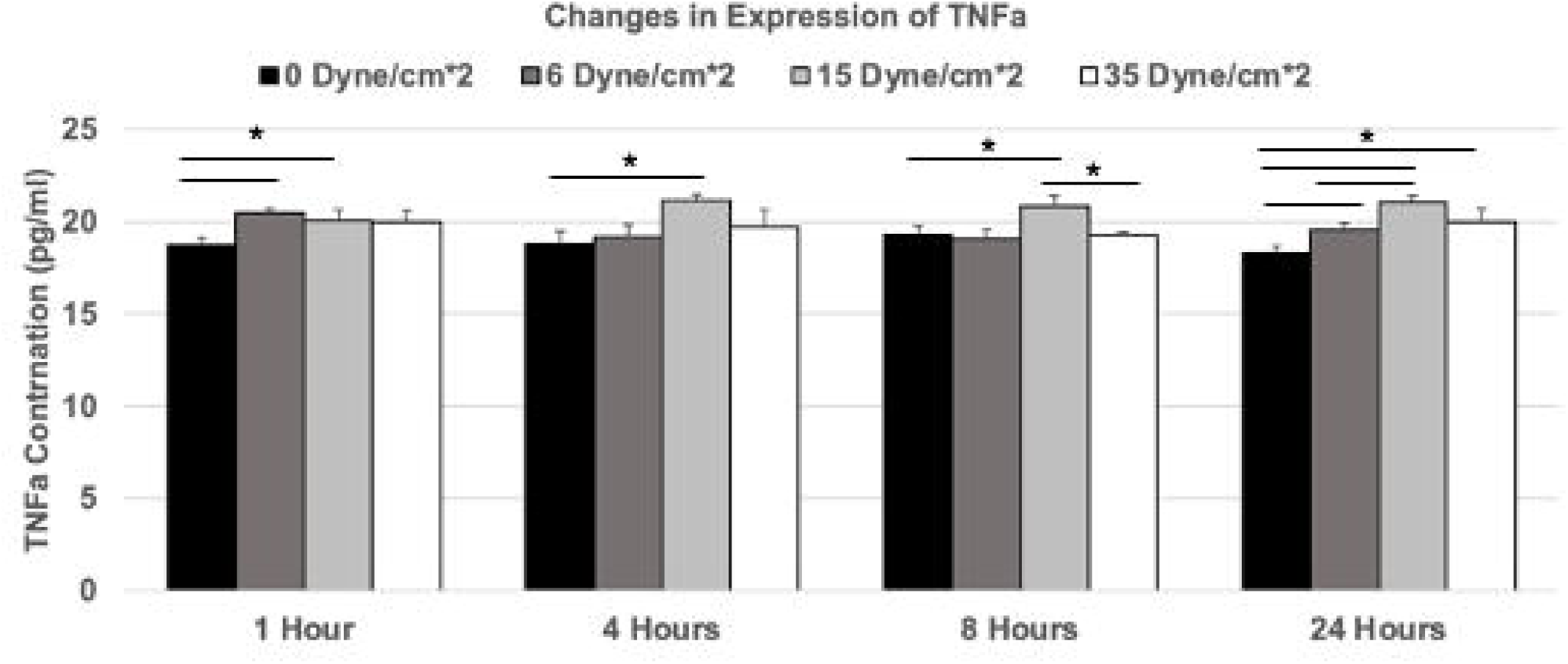
Volcano Plot from Bulk RNA Sequencing. Figure shows individual volcano plots for all comparison groups, followed by a list of the top 10 upregulated and downregulated genes based on significance and the total number of differentially expressed genes for each group.

**Figure 4.**
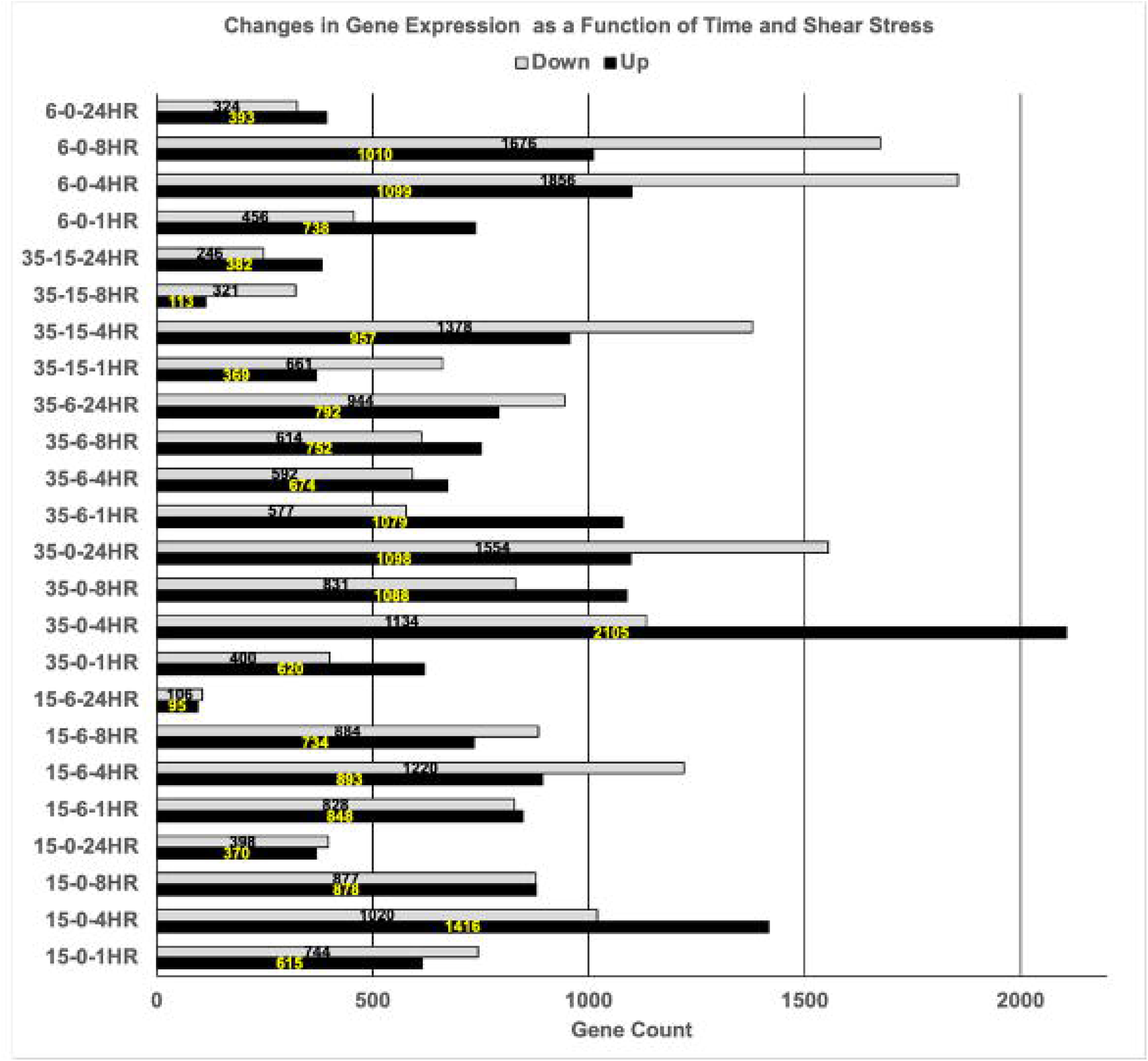
Gene Ontology Pathway Analysis Based on Bulk RNA Sequencing. the top 30 differentially expressed pathways based on significance are shown for all the comparison groups.

**Figure 5.**
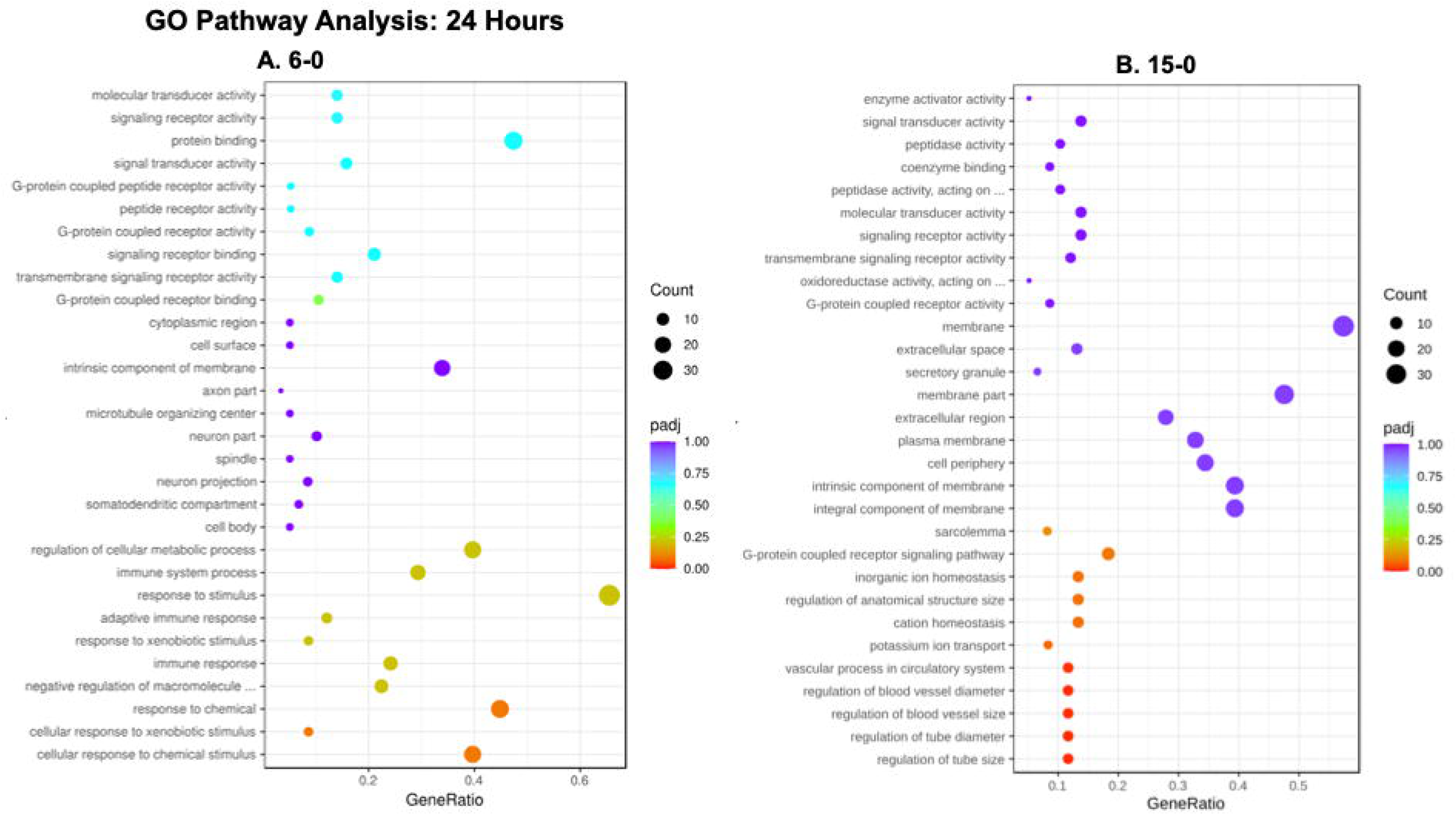
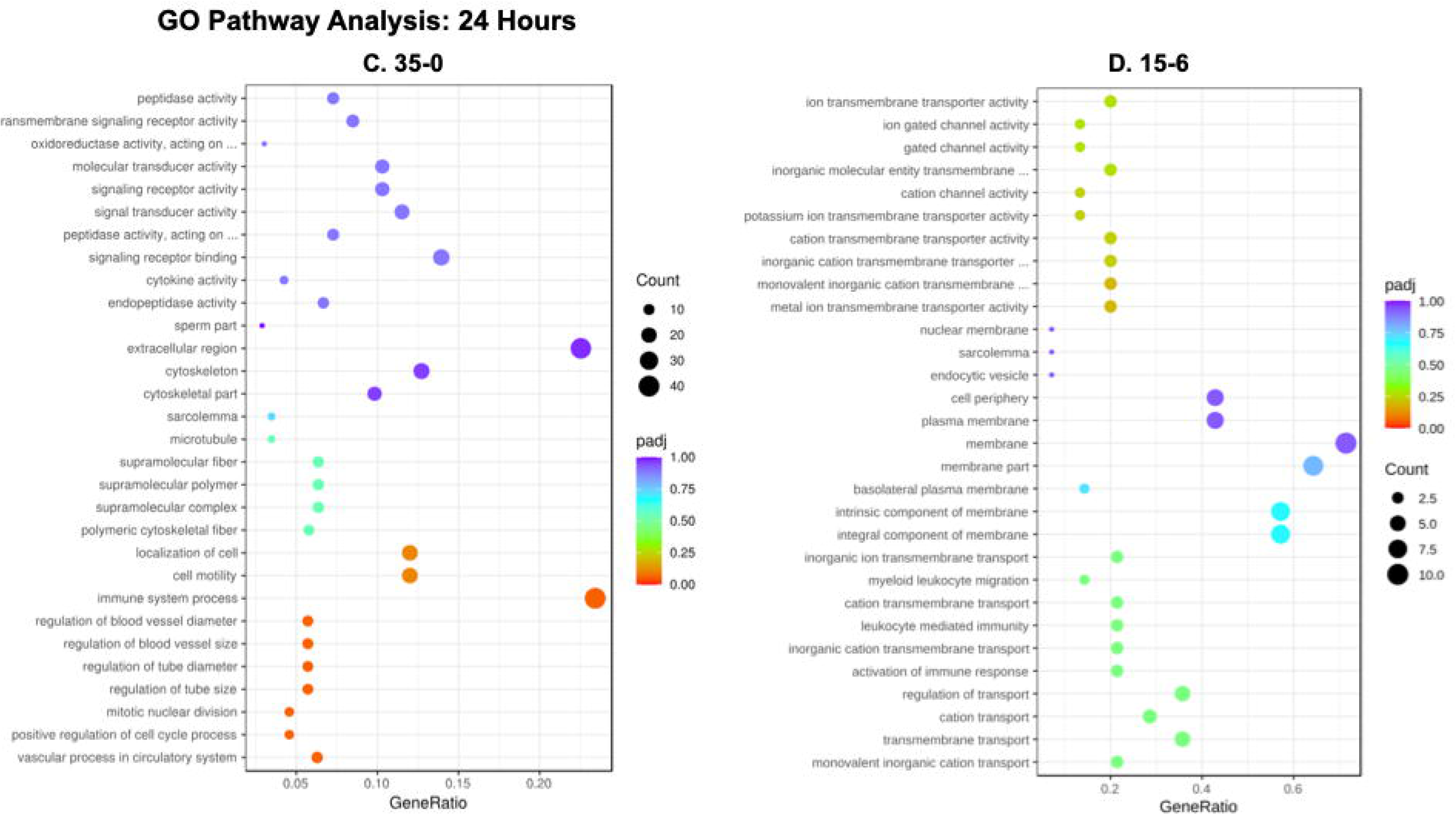
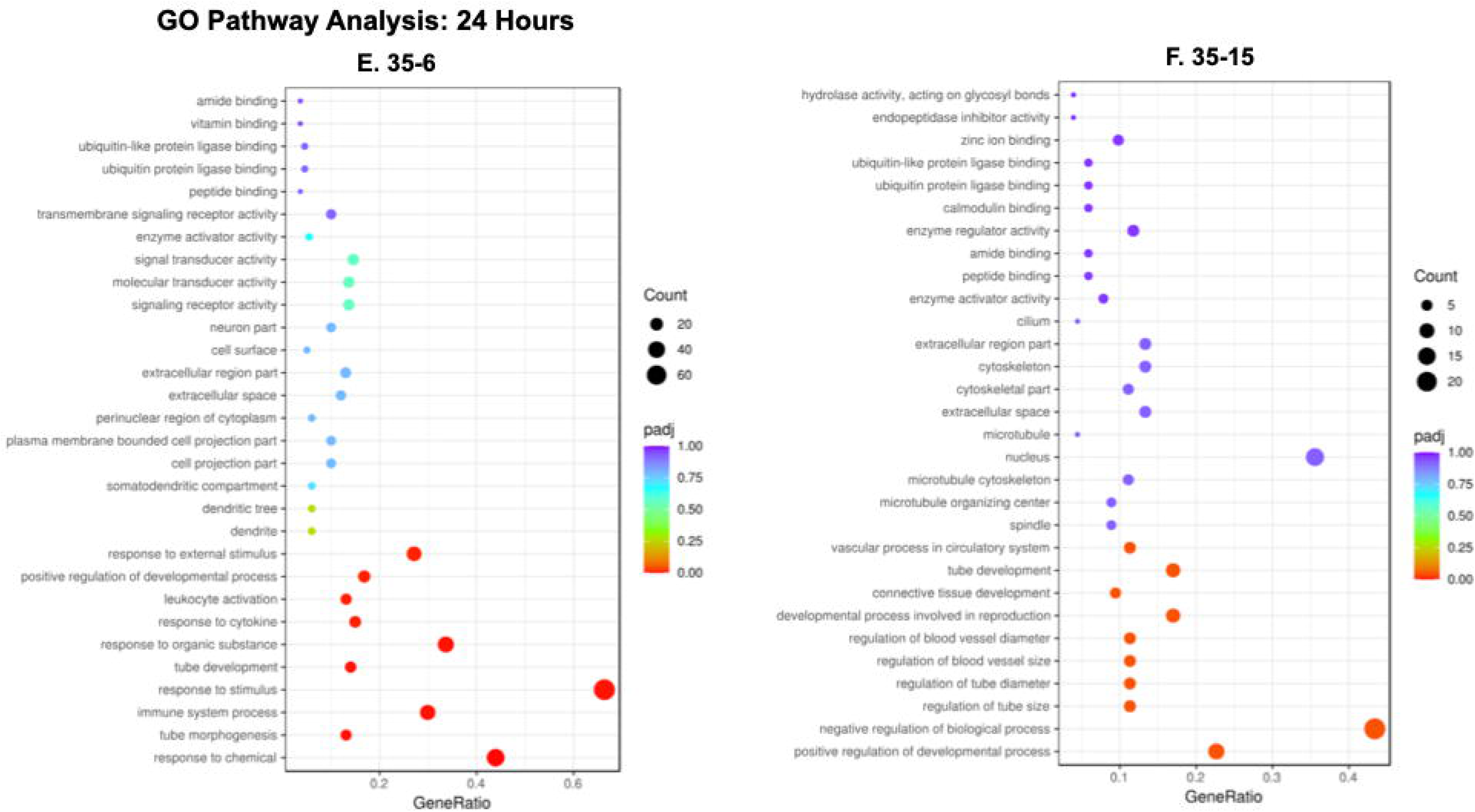
KEGG Pathway Analysis Based on Bulk RNA Sequencing. the top 30 differentially expressed pathways based on significance are shown for all the comparison groups.

**Figures 4 and 5** show the GO analysis and KEGG pathway analysis, respectively. Figure 4 is the Gene Ontology Term Enrichment and named GO enrichment analysis and if there are fewer than 30 terms, all terms are included. These terms are divided into major categories such as biological processes, cell components, molecular functions, and those based on up and down-expressed genes. The X-axis represents the ratio of the number of differential genes associated with a particular GO Term to the total number of differential genes. The Y-axis is the GO Terms. The size of each dot corresponds to the number of genes annotated to a particular GO Term [22]. The data presented in Figure 5 for the KEGG pathway analysis follows a similar scheme. KEGG (Kyoto Encyclopedia of Genes and Genomes) is a collection of manually curated databases containing resources on genomic, biological-pathway and disease information [23]. Pathway enrichment analysis identifies significantly enriched metabolic pathways or signal transduction pathways associated with differentially expressed genes, comparing the whole genome background. Collectively, these results show that inflammatory signaling pathways play an important role in the shear stress response of porcine EECs; in particular, TNFa, RAS and PI3K signaling pathways are significantly upregulated in these cells.

### Downregulation of CD31 expression in porcine EECs attenuates the expression of Inflammatory Cytokines

To determine which siRNA was the best choice for CD31 knockdown in porcine EECs, we obtained 4 custom designed siRNAs from ThermoFisher (#4399665) with gene ID 396941. The immunofluorescence was the positive control to demonstrate transfection in porcine EECs, utilizing BLOCK-iT™ Fluorescent Oligo with FITC (#2013) defined a positive control. As shown in Figure 6A, the positive staining for FITC labeling identified transfected porcine EECs, demonstrating that lipofectamine 3000 transfection was successful and can be used to knockdown CD31. We conducted RT-qPCR to evaluate the CD31 expression in porcine EECs as shown in Figure 6B and determined that No.2 designed siRNA was the best choice for CD31 knockdown since it had the most significant differences among the other groups. Continuously, we utilized the No. 2 designed siRNA with sense sequence 5’CUGCUAAAAUGUAAGAGUAt-3’ and antisense sequence 5’-UACUCUUACAUUUUAGCAGct-3’ to knockdown CD31 in porcine EECs. After CD31 knockdown, the porcine EECs were cultured in the CPBs for 24 hours using all 4 shear stress magnitudes (0, 6, 15, 35 dyne/cm^2^). As was the case before, the conditioned media was collected and the Luminex assay was conducted to determine the concentration of proinflammatory cytokines. As can be seen from Figure 7, the mutant group, defined as CD31 knockdown, had a significantly lower concentration of the following proinflammatory cytokines after 24 hours of culture in the CPBs: GM-CSF, IL-1a, IL-1b, IL-1ra, IL-6, IL-8, IL-18 after 24 hours high shear stress treatment. Also, under low shear stress conditions, there were significant differences between normal and mutant porcine EECs for the following proinflammatory cytokines: GM-CSF, IL-1a, IL-1b, IL-6, IL-12 and IL-18. Collectively, these results serve to demonstrate the role of CD31 in mediating a proinflammatory response in porcine EECs in response to controlled shear stress.

**Figure 6.**
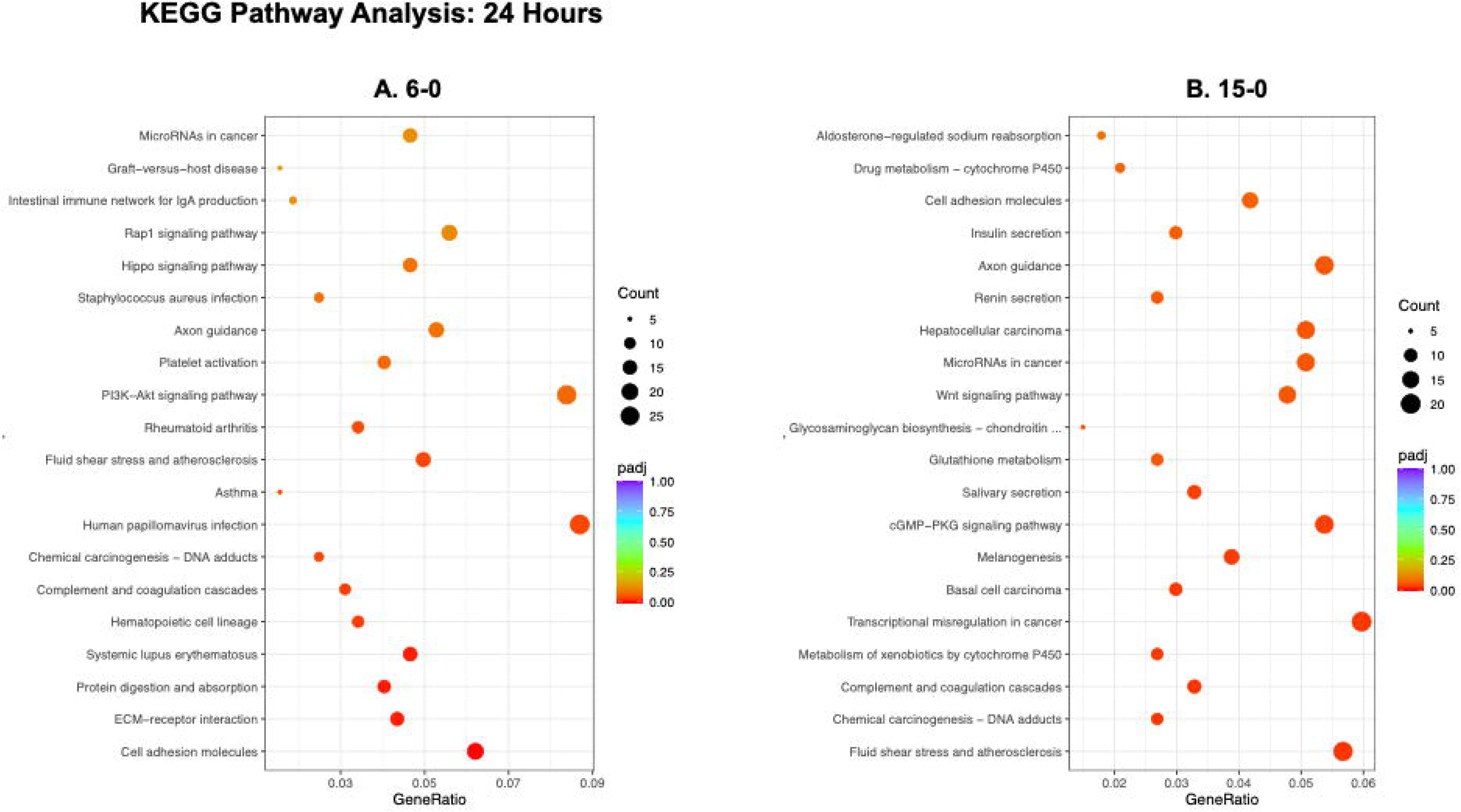
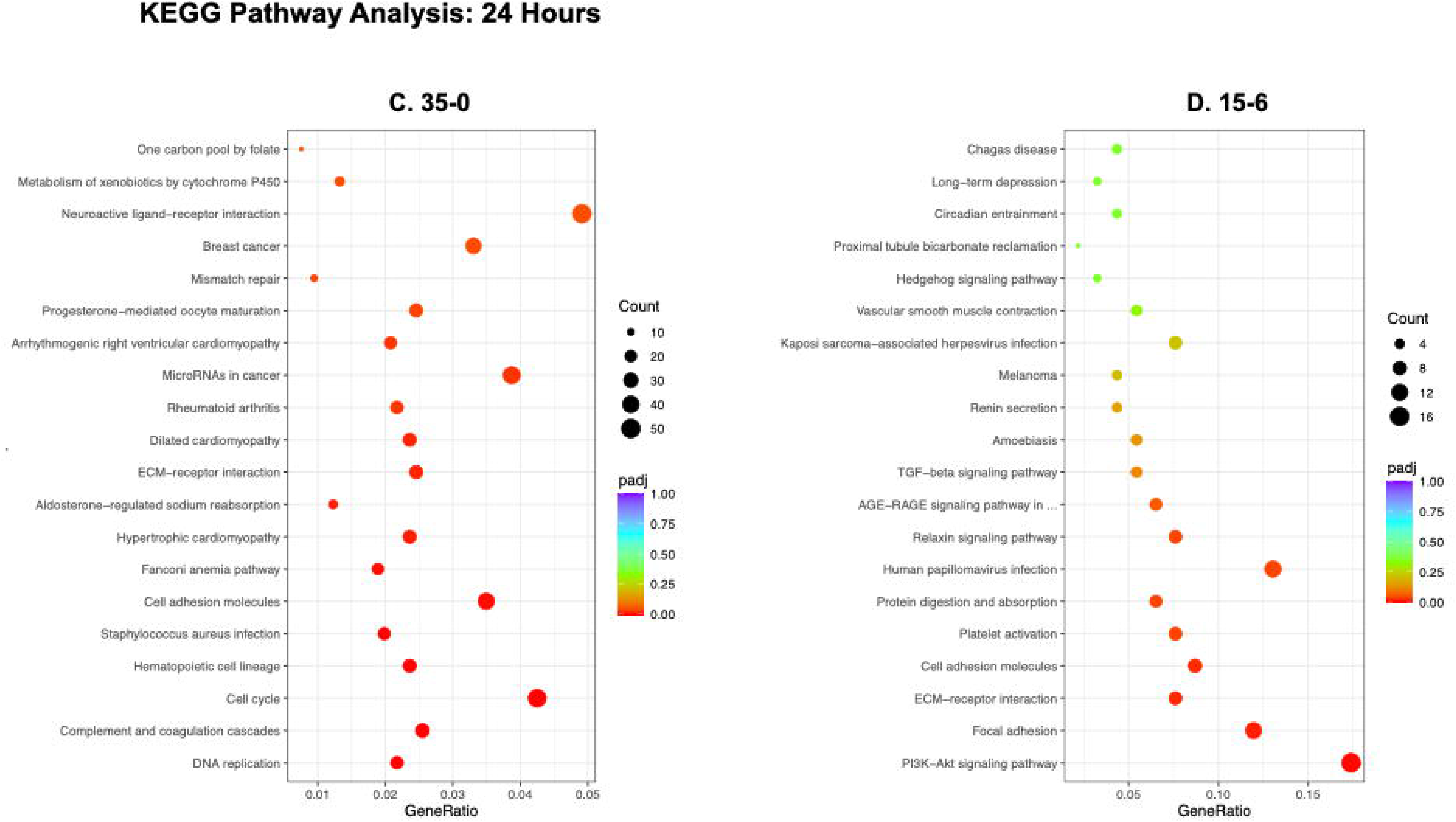
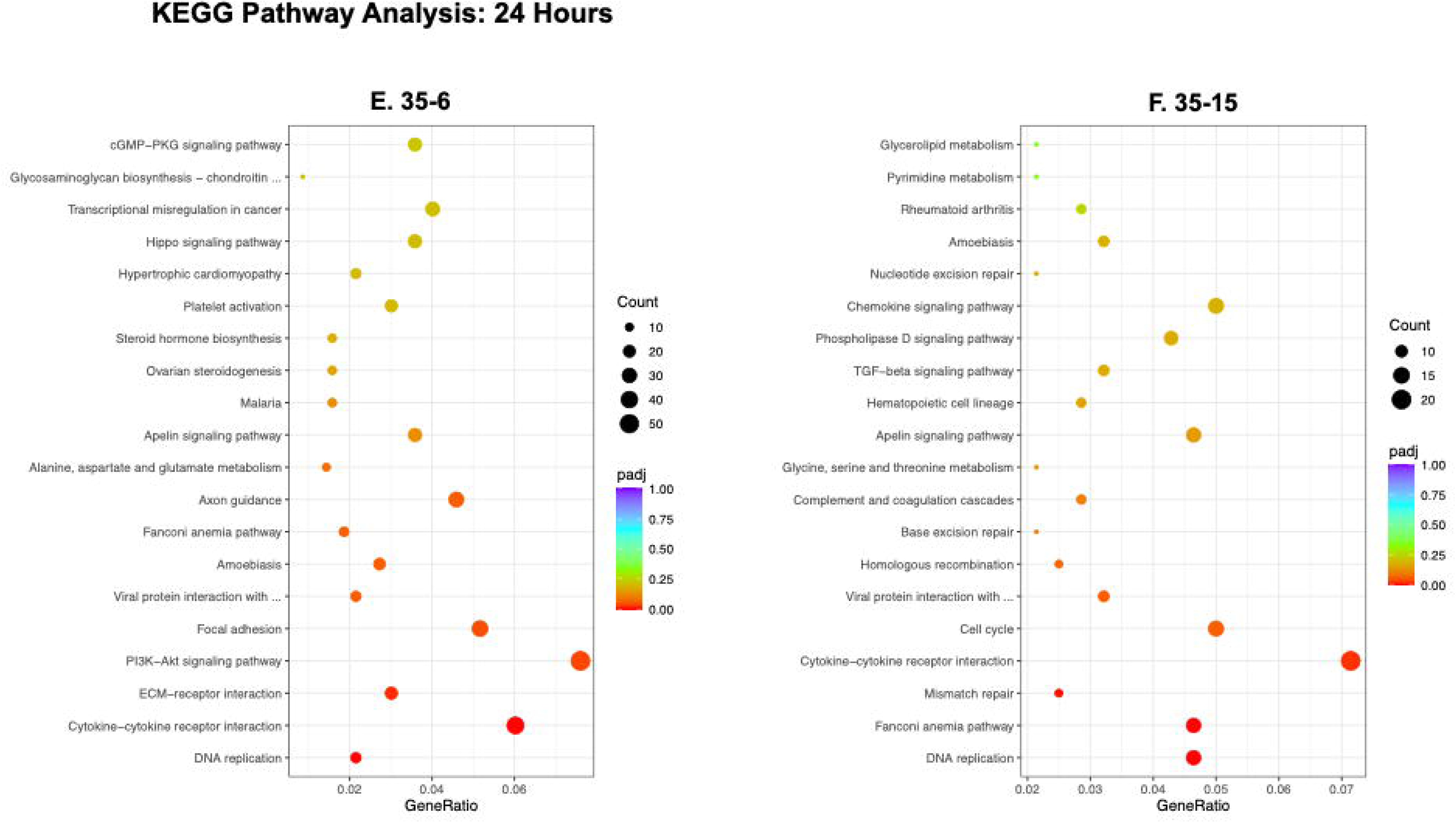
Knockdown of CD31 Expression in EECs.

**Figure 7.**
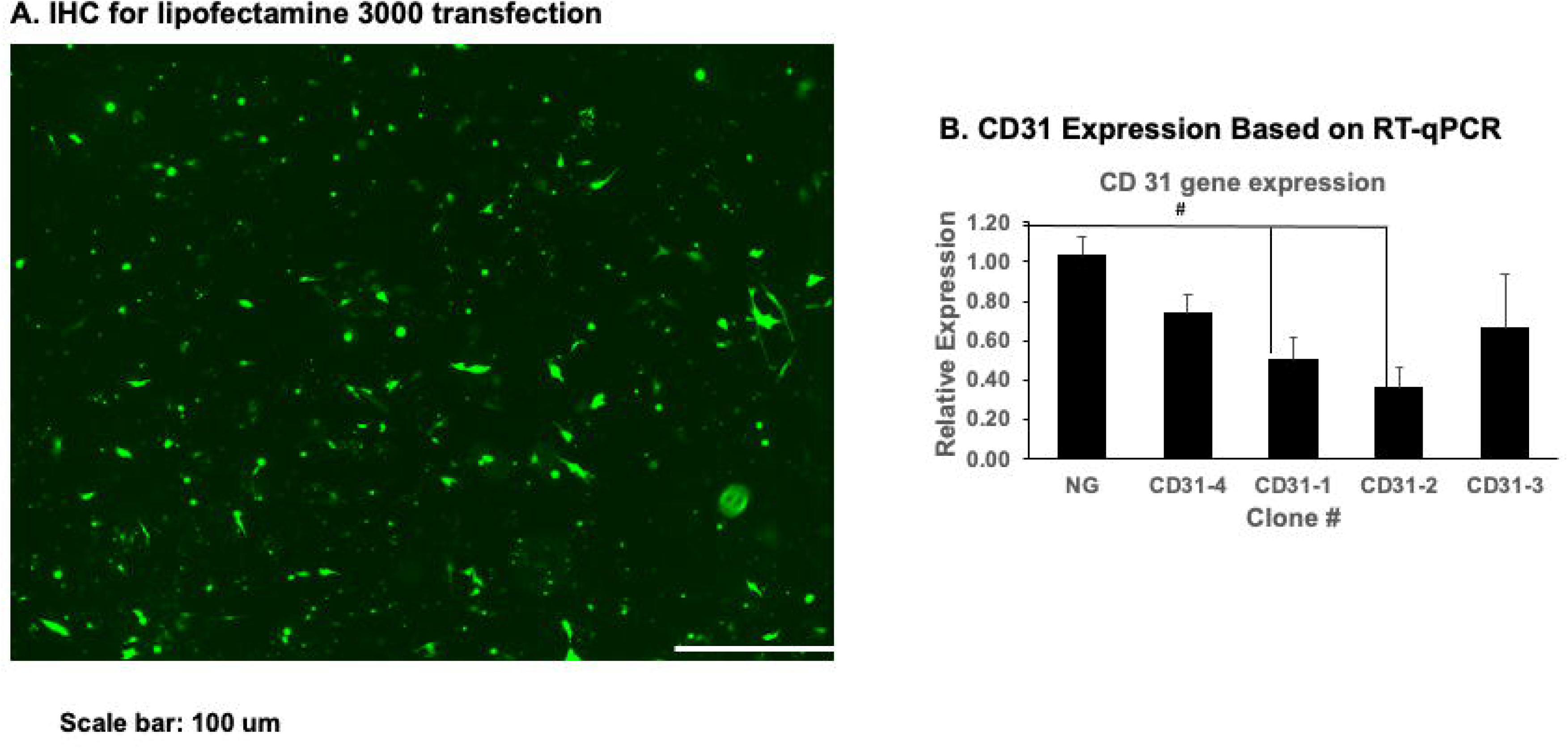
Cytokine Expression after CD31 Knockdown Expression. the Luminex Assay was used to quantify the expression of 13 cytokines for mutant and control EECs - GM-CSF, IL-1b, INFg, IL-1RA, IL-1a, IL-2, IL-4, IL-10, IL-6, IL-12, IL-8, IL-18, TNFa.

**Figure 8.**
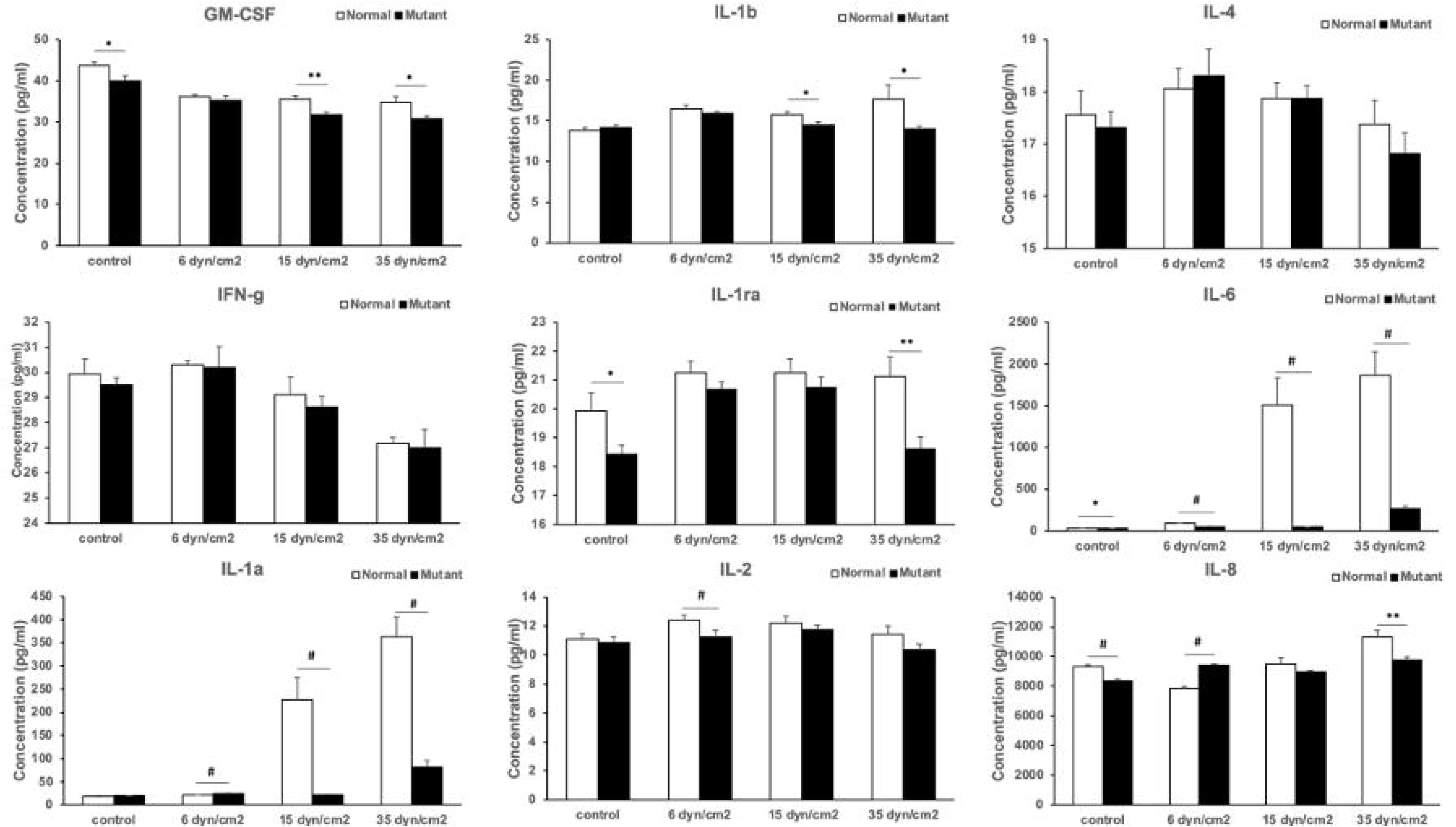

## DISCUSSION

The objective of this study was to develop a model to understand the role of controlled shear stress on EECs within the LVOTs of DSS patients. We hypothesize that EECs sense and respond to changes in the hemodynamic environment in the LVOT and subsequently lead to the formation of the fibrotic membrane. This study was designed to test this hypothesis and characterize the functional response of EECs in response to changes in the shear stress, as a function of time and shear stress magnitude.

During our earlier studies, we developed a computational model that allows us to quantify the magnitude of the shear stress within the LVOT of DSS patients. Based on these computational models, we determined that 15 dyne/cm^2^ and 35 dyne/cm^2^ represented values of shear stress within the LVOT for nonrecurrent and recurrent DSS patients, respectively. We also used two sets of controls, 0 dyne/cm2 representing static culture conditions and 6 dyne/cm^2^, representing physiological shear stress conditions. Furthermore, we utilized porcine EECs as our model to represent EECs within the LVOT, as a source of human cells is not readily available. In our earlier publication, we described the isolation, culture and characterization of EECs from porcine hearts. We further developed a CPB system and demonstrated changes in porcine EECs phenotype based on shear stress exposure. These preliminary studies provided the foundation for the current study.

Our first step was to determine the role of CD31 in mediating the shear stress response of EECs. We accomplished this objective by culturing porcine EECs in different magnitudes of shear stress (0, 6, 15 and 35 dyne/cm^2^) and quantifying the expression of CD31 positive staining (Figure 1). Based on the results of this study, we found an increase in the CD31 positive staining as a function of increasing shear stress magnitude (Figure 1), validating the role of CD31 in mediating the shear stress response of porcine EECs.

After we validated the response of EECs to varying magnitudes of shear stress, our next objective was to quantify the concentration of proinflammatory cytokine released by the EECs in response to shear stress. We accomplished this objective using the multiplex Luminex assay, and quantified the concentration of the following proinflammatory cytokines: GM-CSF, IFN-γ, IL-1a, IL-1b, IL-1RA, IL-2, IL-4, IL-6, IL-8, IL-10, IL-12, IL-18 and TNFa. The rationale for the selection of the cytokines is due to their known roles in inflammatory and briefly described next.

- Granulocyte macrophage colony-stimulating factor (GM-CSF) is a cytokine that stimulates the growth and maturation of myeloid cells, and it plays an important role in promoting the differentiation and survival of dendritic cells [24].
- Interferon gamma (IFN-γ) is an important cytokine involved in initiating and regulating various immune responses. It plays a crucial role in inducing and regulating a variety of immune responses. Intracellular, IFN-γ stimulates the activation of NADPH oxidase and the production of mitochondrial reactive oxygen species (ROS), which contributes to the accumulation of ROS in cells. In addition, IFN-γ triggers activation of JNK (c-Jun N-terminal kinase), ROS, and peroxynitrite, which are key effectors of cell death in IFN-γ responses [25].
- Interleukin1α (IL-1α) and IL-1β are two equally potent inflammatory cytokines that play a key role in activating inflammation. When their signaling becomes out of whack, it can lead to the development of devastating diseases characterized by severe acute or chronic inflammation. These diseases are often accompanied by a marked inflammatory response that can have detrimental effects on the affected individual [26].
- IL-1Ra plays a crucial role in the body’s defense against endotoxin-induced damage. As an anti-inflammatory molecule, IL-1Ra works by reducing the inflammatory effects caused by IL-1. Also, IL-1Ra helps counteract the inflammatory response and mitigate the harmful effects associated with excessive inflammation [27].
- IL-2 is produced by a specific subset of T lymphocytes. It plays a vital role in promoting the growth and enhanced activity of other T and B lymphocytes. In addition, IL-2 plays an important role in the development and maturation of the immune system. By stimulating the proliferation and function of various immune cells, IL-2 contributes to the overall effectiveness of the immune response [28].
- IL-4 and IL-10 are multiple anti-inflammatory cytokines that act primarily by inhibiting the pro-inflammatory environment. IL-4 is produced by a variety of immune cells, including activated T cells, mast cells, basophils, eosinophils, and NKT cells. These cells play a key role in the secretion of IL-4, which contributes to the overall anti-inflammatory response. By suppressing the pro-inflammatory environment, IL-4 and IL-10 help regulate immune response and maintain immune homeostasis [29].
- IL-6 is a multifunctional cytokine that plays a central role in host defense. Its extensive immune and hematopoietic activity, as well as its strong ability to induce an acute phase response. IL-6 plays an important regulatory role in various immune processes, helping the body defend against pathogens and promoting appropriate responses to inflammatory stimuli. In addition, IL-6 plays a key role in regulating hematopoietic production, ensuring that enough immune cells are produced for an effective immune response [30].
- IL-8 is a cytokine known for its pro-inflammatory properties. It plays an important role in the activation of neutrophils, a type of immune cell involved in the body’s defense against infection. IL-8 plays a key role in the onset and progression of a variety of diseases. Its involvement in these diseases contributes to the inflammatory processes, highlighting its relevance in understanding and potentially targeting the underlying mechanisms [31].
- The primary function of IL-12 is to activate T cells and NK cells, thereby increasing INF-γ production, proliferation, and cytotoxic potential. In addition, IL-12 triggers the production of TNF-α, IL-2, IL-3, IL-8, and IL-10. It also stimulates the proliferation of hematopoietic precursors and B cells, further promoting the overall immune response and hematopoietic activity. The multiple roles of IL-12 highlight its critical role in coordinating and amplifying immune responses against pathogens and tumors [32].
- Produced primarily by antigen-presenting cells, IL-18 is a multipotent factor that plays a critical role in the regulation of innate and acquired immune responses. This multifunctional cytokine affects all aspects of the immune system, both the initial innate immune response and the subsequent adaptive immune response. By regulating immune cell activation, cytokine production, and cell communication, IL-18 helps to coordinate the immune responses against pathogens and other immune challenges [33].
- TNF-α (tumor necrosis factor-α) is a potent pro-inflammatory cytokine that plays a crucial role in regulating all aspects of macrophage function. It is a key regulator of inflammatory response and has been implicated in the pathogenesis of several inflammatory and autoimmune diseases. TNF-α acts by affecting immune cell activation, cytokine production, and intercellular communication, ultimately contributing to the inflammatory process in these diseases. Dysregulation of it is signaling can lead to the development and progression of inflammation and autoimmune diseases [34].

Exposing porcine EECs to shear stress resulted in an increase in the release of proinflammatory markers, both a function of time and as a function of shear stress magnitude (Figure 2). While different cytokines spiked at different times and at different magnitudes of shear stress, some very important trends were observed in the expression pattern of IL-8. IL-8 concentration increased with an increase in time and also with an increase in shear stress magnitude, demonstrating the importance of this cytokine in mediating the shear stress response of porcine EECs. Other important cytokines include IL-6, IL-10, IL-12 and IL-18 also showed marked inceases with both time and shear stress magnitude. Collectively, this data served to demonstrate the proinflammatory response of porcine EECs in response to shear stress. This has significant implications for DSS, as it demonstrates the potential role of shear stress mediated inflammatory response of EECs in promoting the formation of fibrotic tissue in DSS patients.

Our next step was to correlate changes in the gene expression profile to changes in shear stress response of porcine EECs. Bulk-RNA sequencing was conducted for porcine EECs as a function of time and as a function of increasing shear stress magnitude and the data presented as differential gene expression (Figure 3), gene ontology analysis (Figure 4) and KEGG pathway analysis (Figure 5).

From the RNA-sequencing data, we choose four significant alterations comparing groups as mentioned above in the result part. From the figures and tables, we found that the potential cell signaling pathways involved with inflammatory signaling pathways and immune cell stimulations. Indeed, the Th1, Th2 and Th17 cell differentiation were activated by the IL-6 and IL-1β, and it also reported that TNF-a also promotes the Th17 cell differentiation [40, 41]. Th1 and Th2 cells play crucial roles in the immune system. Th1 cells are responsible for promoting cellular immune responses, inhibiting macrophage activation, and stimulating B cells to produce antibodies such as IgM and IgG1 [42]. In contrast, Th2 cells stimulate humoral immune responses by promoting B cell proliferation and triggering antibody production, particularly through the release of IL-4. Th2 cells can also induce the differentiation and proliferation of mast cells (via IL-3 and IL-4) as well as eosinophilic leukocytes (via IL-5) [43]. Under normal conditions, the differentiation of Th1/Th2 cells is maintained in a balanced state. However, when this balance is disrupted, it can lead to the development of various diseases Overexpression of Th2 cells can result in inappropriate immune responses, contributing to conditions such as allergies and asthma. Conversely, overexpression of Th1 or Th17 cells can lead to autoimmune diseases like multiple sclerosis [43–44].

Furthermore, the bulk-RNA sequencing show that TNF and PI3K-AKT signaling pathways were highly expressed, which indicated that during the bioreactor treatment the porcine EEC were involved in the inflammation signaling (Figure 6 A-F). The interesting part is that we found the positive control group has promote the neuroactive ligand-receptor interactions and cytokine-cytokine receptor interactions, which mainly involves immune and inflammatory responses [45]. Notably, bioreactor induced the conditioning medium also exhibits common features of the inflammatory in porcine fibroblasts. In summary, these signaling pathways are related to inflammation from above discussions. Also, the immune cells involved in these pathways, such as T cells, neutrophils, macrophages, etc.

Our final validation step was the knockdown of CD31 and subsequent changes in inflammatory cytokines. As shown in Figure 6, we were successful in knocking down the expression of CD31. Subsequently, we repeated the multiplex Luminex assay using the same 13 pro-inflammatory cytokines described before (Figure 7). As shown in Figure 7, knockdown CD31 caused the reduction of inflammatory cytokines. Especially, the mainly significant differences in IL-6 and IL-8. Smooth muscle cells within numerous blood vessels contribute to the production of IL-6, functioning as a pro-inflammatory cytokine [35]. Despite its pro-inflammatory role, IL-6 also serves as an anti-inflammatory myokine by exerting inhibitory effects on TNF-alpha and IL-1, and by activating IL-1ra and IL-10 [36]. Additionally, IL-8 is a chemoattractant cytokine and generated by various tissue and blood cells. In contrast to many other cytokines, IL-8 exhibits distinct target specificity for neutrophils, exerting only weak effects on other blood cells.IL-8 plays a primary role involves attracting and activating neutrophils in inflammatory regions [37]. At both low and high shear stress, there is a significant difference in the expression of IL-1b and GM-CSF. These factors act as triggers to induce a pro-inflammatory response and activate myeloid cells, including neutrophils and macrophages [38]. Furthermore, analysis of IL-12 and IL-18 data suggests that T cells or lymphocytes are stimulated in response to the pro-inflammatory environment, which indicated they try to assist the host to defense the immune response [39].

Subsequently, these findings highlight the interaction of cytokines and immune cells in response to CD31 knockdown, shedding light on the complex dynamics of inflammation and immune regulation. These data were matched with the phenotype of EEC based on cytokine analysis using Luminex data have a lot of points that match with RNA-seq data such as the immune cells involves in the metabolic pathways, several pro-inflammatory cytokines were activated during the bioreactor treatment. Taken together, our studies provide a clear evidence supporting the proinflammatory response of EECs in response to shear stress and demonstrating that CD31 has a role in mediating this response. These results have clear implications for DSS as blocking CD31 expression in DSS patients could provide a potential therapeutic strategy, subsequently reducing the proinflammatory response of EECs and formation of the fibrotic membrane in DSS patients. Future studies will focus on the influence of shear stress released inflammatory cytokines on the growth of the fibrotic membrane in DSS patients. These studies will provide mechanistic insight into DSS and can lead to potential therapeutic strategies.

## ACKNOWLEDGMENTS

This work with supported by an NIH R01 Grant number 5R01HL140305. This work was supported by funding from the Lew and Laura Moorman Family Foundation to SGK. The authors would like to thank financial support from the Division of Congenital Heart Surgery, Texas Children’s Hospital, Houston, TX.

## Funding and/or Conflicts of interests/Competing interests

The authors declare no conflicts of interests.

